# Pathway-Structured Predictive Model for Cancer Survival Prediction: A Two-Stage Approach

**DOI:** 10.1101/043661

**Authors:** Xinyan Zhang, Yan Li, Tomi Akinyemiju, Akinyemi I. Ojesina, Phillip Buckhaults, Nianjun Liu, Bo Xu, Nengjun Yi

## Abstract

Heterogeneity in terms of tumor characteristics, prognosis, and survival among cancer patients has been a persistent problem for many decades. Currently, prognosis and outcome predictions are made based on clinical factors and/or by incorporating molecular profiling data. However, inaccurate prognosis and prediction may result by using only clinical or molecular information directly. One of the main shortcomings of past studies is the failure to incorporate prior biological information into the predictive model, given strong evidence of pathway-based genetic nature of cancer, i.e. the potential for oncogenes to be grouped into pathways based on biological functions such as cell survival, proliferation and metastatic dissemination.

To address this problem, we propose a two-stage procedure to incorporate pathway information into the prognostic modeling using large-scale gene expression data. In the first stage, we fit all predictors within each pathway using penalized Cox model (Lasso, Ridge and Elastic Net) and Bayesian hierarchical Cox model. In the second stage, we combine the cross-validated prognostic scores of all pathways obtained in the first stage as new predictors to build an integrated prognostic model for prediction. We apply the proposed method to analyze breast cancer data from The Cancer Genome Atlas (TCGA), predicting overall survival using clinical data and gene expression profiling. The data includes ~20000 genes mapped into 109 pathways for 505 patients. The results show that the proposed approach not only improves survival prediction compared with the alternative analysis that ignores the pathway information, but also identifies significant biological pathways.

## INTRODUCTION

Over the past three decades, remarkable improvement has been achieved in cancer treatment in the United States, with the annual death rate from cancer declining 1.4% for women, 1.8% for men, and 2.3% for children ages 0-10 years from 2002 to 2011 (EDWARDS *et al*. 2014). However, the problem that has persisted in cancer treatment is the heterogeneity of prognostic prediction across patients (BARILLOT 2013). This heterogeneity is, for the most part, genetically determined and rooted in the molecular profile of patients. A Precision medicine initiative has been introduced by White House to expand cancer genomics research as a short-term goal to develop better prevention and treatment methods for more cancers (COLLINS and VARMUS 2015). Recent high-throughput technologies can easily and robustly generated large-scale molecular profiling data, including genomic, epi-genomic, transcriptomic, and proteomic markers offers extraordinary opportunities to integrate clinical and genomic data in prediction models, improving understanding of inter-individual differences that may be critical for the application of precision medicine strategies (BARILLOT 2013; COLLINS and VARMUS 2015).

The recent development of molecular signatures to predict recurrence of breast, colon and prostate cancers are notable and clinically useful but may not be sufficient to achieve the goals of precision medicine (MOOK *et al*. 2007; POHL and LENZ 2008; ENG *et al*. 2013). Gene signatures across different studies with very few overlapped genes can have similar prediction results which suggests there is some underlying mechanism (VAN DE VIJVER *et al*. 2002; WANG *et al*. 2005; SOTIRIOU and PICCART 2007). According to the analysis of 24 pancreatic tumors by JONES et al (2008), altered genes varied greatly across tumors but the pathways with the altered genes remain largely the same, which indicates that the statistical methods focusing on individual genes may be underpowered. It has also been revealed that the genetic nature of cancer is pathway-based, that is, oncogenes can be grouped into pathways based on biological functions such as cell survival, proliferation and metastatic dissemination (BARILLOT 2013; HUANG *et al*. 2014). Based on the observation that multiple genes in the same biological processes appear to be dysfunctional regardless of cancer type, gene pathways information is likely a more robust biological phenomenon (BILD *et al*. 2006). Various public databases (e.g., KEGG) can be accessed online or in R package to provide the biological information about pathways which may provide valuable improvement to prognosis and prediction (KANEHISA and GOTO 2000). Thus methods incorporating higher-order information of functional units in cancer, i.e., pathways, have been the focus of recent investigations (JONES 2008; JONES *et al*. 2008; LEE *et al*. 2008; REYAL *et al*. 2008; ABRAHAM *et al*. 2010; TESCHENDORFF *et al*. 2010; ENG *et al*. 2013; HUANG *et al*. 2014). Among those previous studies, Abraham et al. adopted a gene set statistic to provide stability of prognostic signatures instead of individual genes (ABRAHAM *et al*. 2010). Huang et al. converted the gene matrix to a pathway matrix through “principal curve”, similar to principal components analysis (HUANG *et al*. 2014). Both of these two methods did not incorporate outcome when generating the pathways scores from the individual genes. Other sophisticated statistical methods have been developed for variable selection with grouped predictors or pathways using an “all-in-all-out” idea, meaning that when one predictor in a group is chosen, then all variables in that group are chosen (PARK *et al*. 2007; WEI and LI 2007; JONES 2008). Other methods that could address the above shortcomings have also been developed, however, leading to increased computational complexity and potentially instability of models when the number of predictors is large (HUANG *et al*. 2009; ZHOU 2010; ENG *et al*. 2013). ENG et al. (2013) proposed a method to reduce the computational complexity by incorporating a binary outcome to stand for decreased or increased risk score in each pathway which inferred potentially loss of information.

In this article, to address some of the above shortcomings, we propose a two-stage procedure to incorporate pathway information into the prognostic models using large-scale gene expression data. In the first stage, we fit all predictors within each pathway using penalized Cox model (Lasso, Ridge and Elastic Net) and Bayesian hierarchical Cox model. In the second stage, we combine the cross-validated prognostic scores of all pathways obtained in the first stage as new predictors to build an integrated super prognostic model for prediction. We used the proposed method to analyze a breast cancer data set from The Cancer Genome Atlas (TCGA) project for predicting overall survival using gene expression profiling.

Breast cancer is the second most commonly diagnosed malignancy after skin cancer in women (HUANG *et al*. 2014). It is estimated to be the third leading cause of cancer death after lung cancer and rectal/colon cancer in 2015 (NCI 2015). It is widely understood that breast cancer can be categorized into four clinical subtypes: Luminal A, Luminal B, Triple Negative/Basal like and Her2 and the survival/metastasis outcomes differ significantly among these four subtypes (CAREY *et al*. 2006; O'BRIEN *et al*. 2010; HAQUE *et al*. 2012). However, it is increasingly being realized that using only the clinical subtypes cannot discriminate breast cancers patients, and that better prediction of prognosis is needed. The breast cancer data set from TCGA includes ~20000 genes mapped into 109 pathways for 505 patients. The results show that the proposed approach not only improves survival prediction compared with the alternative analysis that ignores the pathway information, but also identifies significant biological pathways.

## METHODS

### Cox proportional hazards models

Cox regression is the commonly used method for analyzing censored survival data (VAN HOUWELINGGEN and PUTTER 2012), for which the hazard function of survival time *T* takes the form:

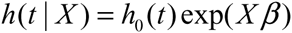

where *h*_0_(*t*) is the baseline hazard function, *X* and *β* are the vectors of predictors and coefficients, respectively, and *Xβ* is the linear predictor or called the prognostic index. The coefficients *β* are estimated by maximizing the partial log-likelihood:

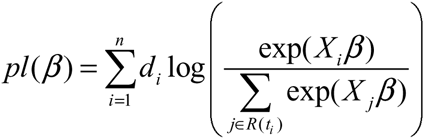

where the censoring indicator *d*_*i*_ takes 1 if the observed survival time *t*_*i*_ for individual *i* is uncensored and 0 if it is censored, and *R*(*t*_*i*_) is the risk set at time *t*_*i*_. For molecular data, the number of coefficients is much larger than the number of individuals and/or covariates are usually highly correlated, where Cox regression is not directly applicable.

### Ridge, lasso and elastic-net Cox models

The elastic net is a widely used penalization approach to handle high-dimensional models, which adds the elastic-net penalty to the log-likelihood function and estimates the parameters *β* by maximizing the penalized log-likelihood(ZOU and HASTIE 2005; HASTIE *et al*. 2009; FRIEDMAN *et al*. 2010; SIMON *et al*. 2011; HASTIE *et al*. 2015). For the Cox models described above, we estimate the parameters *β* by maximizing the penalized partial log-likelihood:

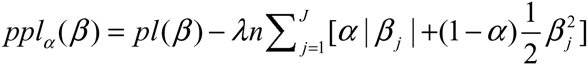

where *α* (0 ≤ *α* ≤ 1) is a predetermined elastic-net parameter, *λ* (*λ* ≥ 0) is a penalty parameter, and *pl*(*β*) is the partial log-likelihood of the Cox model. The penalty parameter *λ* controls the overall strength of penalty and the size of the coefficients; for a small *λ*, many coefficients can be large, and for a large *λ*, many coefficients will be shrunk towards zero. The elastic net includes the lasso (*α* = 1) and ridge Cox regression (*α* = 0) as special cases (TIBSHIRANI 1997; GUI and LI 2005; VAN HOUWELINGEN *et al*. 2006; SIMON *et al*. 2011; VAN HOUWELINGGEN and PUTTER 2012).

The ridge, lasso and elastic net Cox models can be fitted by the cyclic coordinate descent algorithm, which successively optimizes the penalized log-likelihood over each parameter with others fixed and cycles repeatedly until convergence. The cyclic coordinate descent algorithm has been implemented in the R package glmnet. The package glmnet can quickly fit the elastic-net Cox models over a grid of values of *λ* covering the entire range, giving a sequence of models for users to choose from. Cross-validation is the most widely used method to select an optimal value *λ* (e.g., an optimal Cox model) that gives minimum cross-validated error.

#### Bayesian hierarchical Cox model

hierarchical model is an efficient approach to handling high-dimensional data, where the regression coefficients are themselves modeled (GELMAN and HILL 2007; GELMAN *et al*. 2014). Hierarchical models are more easily interpreted and handled in the Bayesian framework where the distribution of the coefficient is the prior distribution, and statistical inference is based on the posterior estimation. The commonly used prior is the double-exponential (or Laplace) prior distribution (PARK and CASELLA 2008; YI and XU 2008; YI and MA 2012):

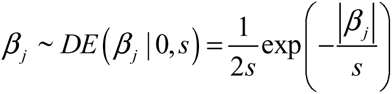

where the scale *s* is shrinkage parameter and controls the amount of shrinkage; a smaller scale s induces stronger shrinkage and thus forces the estimates of *β*_*j*_ towards the prior mean zero. For the hierarchical Cox model with the double-exponential prior, the log posterior distribution of the parameters can be expressed as

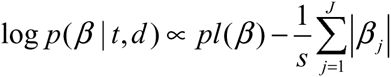

We fit the hierarchical Cox model by finding the posterior modes of the parameters, i.e., estimating the parameters by maximizing the log posterior distribution. We have developed an algorithm for fitting the hierarchical Cox model by incorporating an EM procedure into the usual Newton-Raphson algorithm for fitting classical Cox models. Our algorithm has been implemented in R package BhGLM (http://www.ssg.uab.edu/bhglm/).

### Relation of hierarchical Cox model to the Lasso

The Lasso is equivalent to hierarchical Cox model with the double-exponential prior 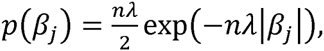 if one estimates the mode of the posterior distribution. With the double-exponential prior, therefore, the relation between hyper-parameter inverse scale *s* and penalty tuning parameter *λ* of Lasso is *s* = 1/(n*λ*).

### Optimizing the penalty by cross-validation and selection of inverse scale

The penalized Cox model estimate of the previous section depends heavily on the penalty tuning parameter *λ*. If *λ* is taken too small, the model boils down to the conventional model virtually with no penalty and the solution still degenerates. If *λ* is taken too large, all the coefficients are estimated to virtually zero. The tuning parameter is estimated using 10-fold cross-validation over 10 repeats by maximizing the cross-validated partial likelihood (CVPL) *CV*(*λ*) (VAN HOUWELINGEN *et al*. 2006; SIMON *et al*. 2011):

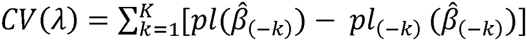

*β̂*(–*k*)is the estimate of β from all the data except the *k*-th part, *pl*(*β̂*(_–*k*_)) is the partial likelihood of all the data points and *pl*(_–*k*_)(*β̂*(_–*k*)_) is the partial likelihood excluding part *k* of the data. By subtracting the log-partial likelihood evaluated on the non-left out data from that evaluated on the full data, we can make efficient use of the death times of the left out data in relation to the death times of all the data. We choose the *λ* value which maximizes *CV*(*λ*). Hyper-parameter inverse scale in hierarchical Cox model is estimated using conversion from estimated tuning parameter *λ* of Lasso with *s* = 1/(*nλ*).

### Two-Stage Approach for Pathway Integration

Intuitively, we can simultaneously fit all the genes in one model. In some previous studies, penalized Cox model (Lasso, Ridge and Elastic Net) have been used to analyze genomic data with this gene-based single model approach (RAPPAPORT 2007; BOVELSTAD *et al*. 2009; JACOB 2009; ZHANG *et al*. 2013; YUAN *et al*. 2014; ZHAO *et al*. 2014). However, due to high-dimension of genomic data, fitting one model including all genes can lead to instability of predictive model, and may result in bad prediction performance with the increased model complexity.

Here we used an two-stage procedure for combining multiple pathways to build a prediction model, inspired by the super learner of VAN DER LAAN et al. (2007) (VAN DER LAAN *et al*. 2007; VAN HOUWELINGEN and PUTTER 2011). The two-stage procedure for building the prognostic model by combining multiple biological pathways is presented in Figure 1.

**Figure 1.**
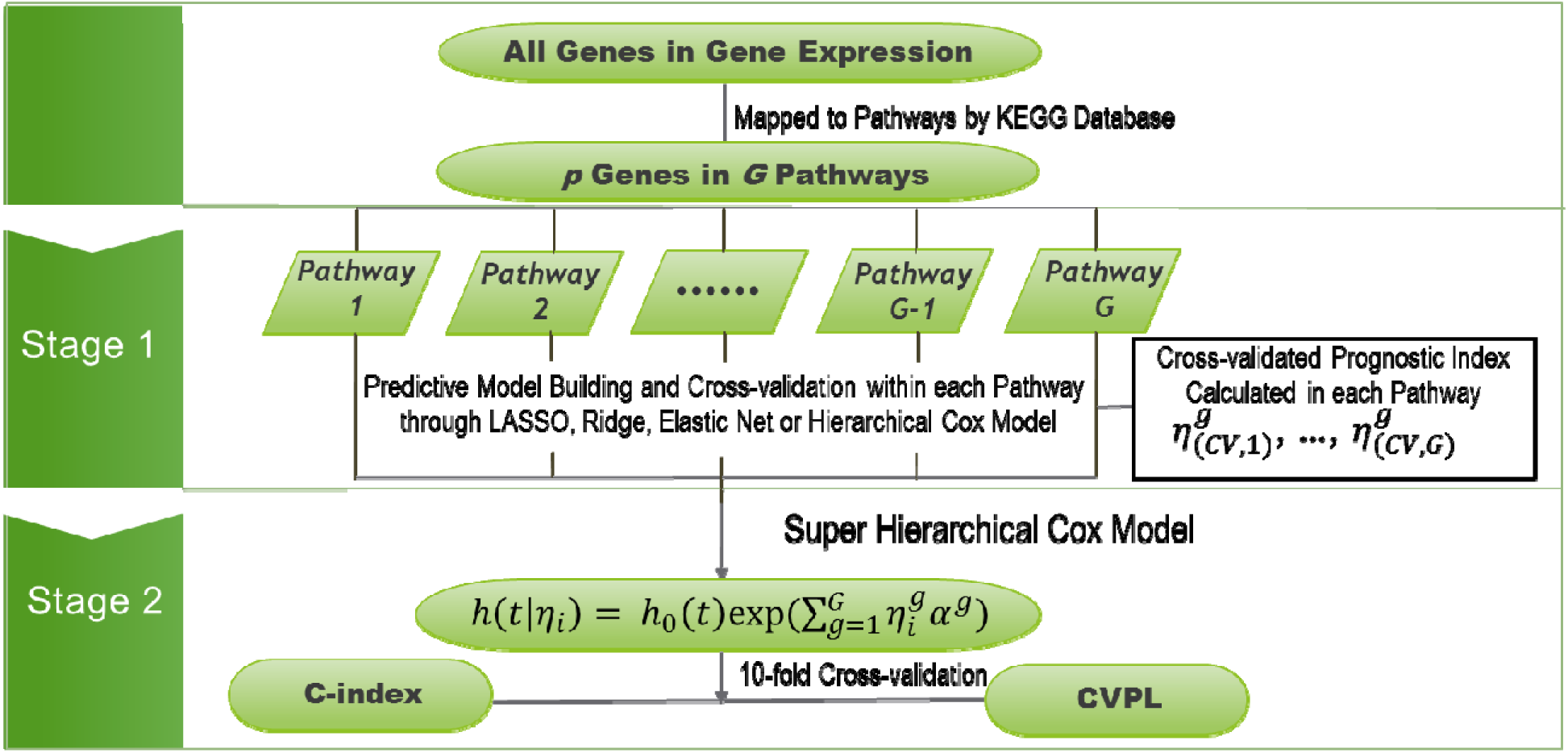
Flowchart of the two-stage prognostic model.

Using the notations from Figure 1, gene expression could be divided into *G* groups (e.g., pathways), *G*_*g*_:*g* = 1, …, *G*, with the *g-th* group *G*_*g*_ containing *J*_*g*_ variables. Overlapping is common in this analysis, that is, a gene could belong to multiple functional pathways. In the first stage, we fit all the predictors within *g-th* group (pathway) using the hierarchical Cox model or penalized Cox regression. For *g-th* pathway, we have hazard function: 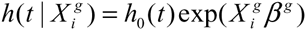 and obtain the estimate of prognostic score, 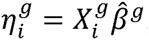, for each individual. To prevent over-fitting, instead of calculating the prognostic indices directly, we estimated cross-validated prognostic scores using leave-one-out cross-validation (LOOCV). The prognostic scores were calculated for *i-th* testing set using the parameters *β̂*(*g*^*−i*^) estimated from (*i*-1) training set. That is,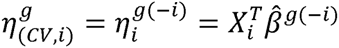. In the second stage, we combine the cross-validated prognostic scores from all pathways as new predictors to build a super prognostic model for prediction: 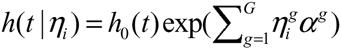 10-fold cross-validation over 10 repeats was used to evaluate the predictive performance of the super prognostic model.

### Evaluating the predictive performance

To assess the prognostic utility of the fitted model, we need to evaluate the quality of the fitted model and its predictive value. There are several ways to measure the performance of a Cox model (STEYERBERG 2009; VAN HOUWELINGGEN and PUTTER 2012): 1) **Concordance Index (C-index)** (HARRELL *et al*. 1996): traditional measurement to determine the concordance between the observed survival times and predicted survival times. The performance is better when the C-index is greater; 2) **CVPL:** as mentioned above as *CV*(*λ*), is a general measurement of model quality and prediction; 3) **Prediction Error**: Measuring prediction error is an important way to evaluate predictive performance of a survival model. The most popular measure of prediction error is the Brier score, which is defined as (VAN HOUWELINGGEN and PUTTER 2012): *Brier*(*y*, *S*(*t*_0_| *x*)) = (*y* – *S*(*t*_0_| *x*))^2^, where *S*(*t*_0_ | *x*) is the estimated survival probability of an individual beyond t_0_ given the predictor *x*; 4) **Pre-validated Kaplan Meier Analysis**: We transform the continuous cross-validated prognostic score 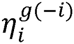 from the super prognostic model into a categorical factor based on the median of 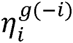 and give the Kaplan Meier plot and log-rank test by comparing the 2 groups splitting by median. This allows us to compare statistical significance and prediction performance between models.

## APPLICATION TO TCGA BREAST CANCER DATA

### Data Collection

1071 tumor samples of breast cancer were selected from TCGA breast cancer project. All data including clinical information, microarray mRNA gene expression were downloaded from TCGA as of February 2015 using the *TCGA-Assembler* (ZHU et *al*. 2014). Overall survival (OS) is the outcome of interest. We downloaded and analyzed the processed level 3 (log2 lowess normalized (cy5/cy3) collapsed by gene symbol) gene expression data. It represents up-regulation or down-regulation of a gene compared relative to the reference population (ZHAO *et al*. 2014).

### Data Preprocessing

54 samples were removed for missing or zero overall survival time. 17815 features across 533 samples were profiled for gene expression, which includes a total of 1571 missing observations. Simple imputation with mean values across samples was adopted to fill the missing values. For further analysis, only 505 samples were kept for whom survival time and gene expression were both available. Among these 505 patients, only 65 were dead and thus the event rate was 12.9%.

### Pathway Analysis

To construct the pathways, we used genome annotation tools, KEGG (KANEHISA and GOTO 2000), to map genes to pathways. After mapping gene symbols to Entrez ids, 17252 probes were kept. We mapped all the probes to KEGG pathways using the R annotation package *RDAVIDWebService* (FRESNO and FERNANDEZ 2013). 3181 probes were mapped to 109 pathways.

### Two-Stage Approach for integrating pathways

#### Separately analyzing gene expression data in each pathway

We compared Lasso, Ridge, Elastic Net (*α* = 0.5) regression and hierarchical Cox models to build predictive model and calculate the LOOCV prognostic scores in each pathway in the first stage. The tuning parameters in Lasso, Ridge and Elastic Net regressions were estimated by 10-fold cross-validation over 10 repeats. We obtained the estimated tuning parameter *λ* from Lasso in each pathway. We used *s*_1_ = 1/*nλ*, *s*_2_ = 1/*nλ* + 0.03 and *s*_3_ = 0.08 as the scales of the double-exponential prior for hierarchical Cox model. The table Appendix 1 shows the cross-validated partial likelihood (CVPL) and C-index for each pathway from LOOCV for the hierarchical Cox models (*s*_1_ = 1/*nλ*, *s*_2_ = 1/*nλ* + 0.03 and *s*_3_ = 0.08, respectively), Lasso, Ridge and Elastic Net Cox regression, respectively.

#### Combining all pathways for survival prediction

In the second stage, only pathways with C-index greater than 0.5 were kept to build a super prognostic model. A super prognostic model was built with the cross-validated prognostic scores of all filtered pathways estimated in the first stage as new predictors. We used hierarchical Cox model with double exponential prior to fit this super prognostic model. 10-fold cross-validation over 10 repeats was also carried out to validate the second stage super prognostic models. To select the prior scales for hierarchical Cox models, we calculated CVPL from 10-fold cross-validation for different prior scales (0.08, 0.10, 0.12, 0.14, 0.16, 0.18, 0.20, and 0.22) and the scale with highest CVPL was chosen. Table 1 shows CVPL, C-index and corresponding prior scales for hierarchical Cox models for all two-stage Lasso-hierarchical Cox Model, two-stage Ridge-hierarchical Cox Model, two-stage Elastic Net-hierarchical Cox Model and three two-stage hierarchical-hierarchical Cox Model (*s*_1_ = 1/*nλ*, *s*_2_ = 1/*nλ* + 0.03 and *s*_3_ = 0.08, respectively). CVPL for two-stage hierarchical-hierarchical Cox Model (*s*_1_ = 1/*nλ* and *s*_2_ = 1/*nλ* + 0.03) and two-stage Lasso-hierarchical Cox Model are -337.170 (7.187), -333.358 (1.971) and -340.058 (4.215). C-index for two-stage hierarchical-hierarchical Cox Model (*s*_1_ = 1/*nλ* and *s*_2_ = 1/*nλ* + 0.03) and two-stage Lasso-hierarchical Cox Model are 0.760 (0.015), 0.748 (0.013) and 0.725 (0.014).

**Table 1.** Prediction Performance Comparison between Different Two-stage Models.

| **Model** | **Number of Pathways** | **CVPL** | **C-index** | **Prior Scale Second Stage hierarchical Model** |
| --- | --- | --- | --- | --- |
| Two-stage Lasso-hierarchical Cox Model | 75 | -340.058 (4.215) | 0.725 (0.014) | 0.20 |
| Two-stage Ridge-hierarchical Cox Model | 88 | -353.444 (2.373) | 0.640 (0.021) | 0.10 |
| Two-stage Elastic Net-hierarchical Cox Model<br>($\alpha=0.5$) | 74 | -340.893 (2.435) | 0.711 (0.012) | 0.16 |
| Two-stage hierarchical-hierarchical Cox Model<br>($s_1 = 1/n\lambda$) | 77 | -337.170 (7.187) | 0.760 (0.015) | 0.26 |
| Two-stage hierarchical-hierarchical Cox Model<br>($s_2 = 1/n\lambda + 0.03$) | 77 | -333.358 (1.971) | 0.748 (0.013) | 0.14 |
| Two-stage hierarchical-hierarchical Cox Model<br>($s_3 = 0.08$) | 70 | -347.459 (2.796) | 0.692 (0.014) | 0.16 |

### Prediction Performance Comparison between Single Model Analysis and Two-stage Approach

To compare the predictive performance with the two-stage approach, a joint Lasso and joint hierarchical Cox model were built which simultaneously fit all 3181 genes from 109 pathways. The tuning parameter *λ* in joint Lasso was estimated to be 0.057 by 10-fold cross-validation over 10 repeats. We used s = 1/*nλ* =0.035 as the scale of the double-exponential prior for joint hierarchical Cox model. 10-fold cross-validation over 10 repeats was used to validate joint Lasso and joint hierarchical Cox model. CVPL and C-index for both joint Lasso and joint hierarchical Cox model are presented in Table 2. CVPL and C-index for joint Lasso are -364.845 (0.949) and 0.507 (0.023), respectively. CVPL and C-index for joint hierarchical Cox model are -363.554 (0.626) and 0.572 (0.023), respectively. Both of them had worse prediction performance than our proposed two-stage combination method.

**Table 2.**
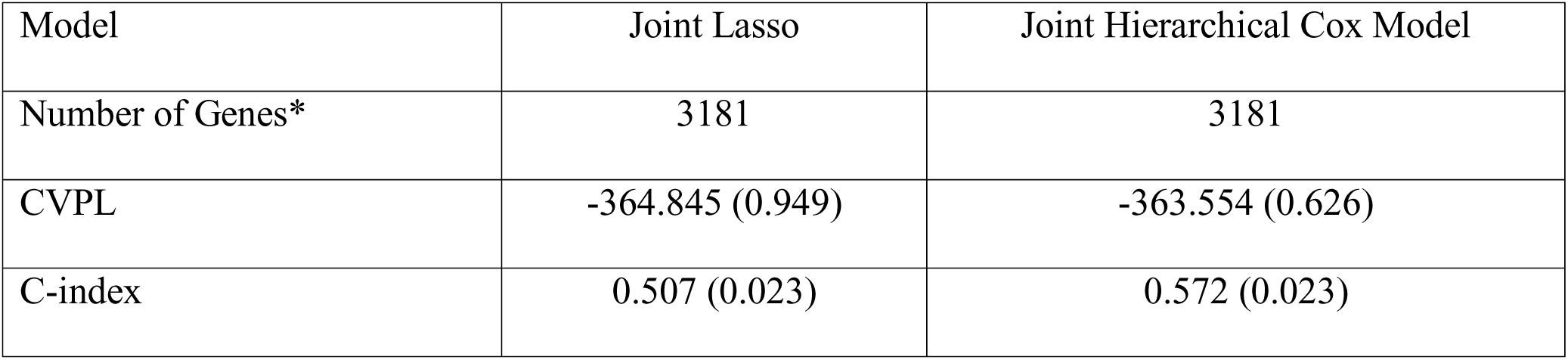
Prediction performance comparison between two single models.

The predictive performance of the models was also assessed by Brier scores. Figure 1 shows the Brier prediction errors for two-stage hierarchical-hierarchical Cox Model (*s*_1_ = 1/*nλ*, *s*_2_ = 1/*nλ* +0.03), two-stage Lasso-hierarchical Cox Model, and two-stage Ridge-hierarchical Cox Model, the hierarchical Cox fitted pathway with the best predictive performance and joint Lasso. Our two-stage models all had significantly improvement in reducing prediction error compared with best hierarchical Cox fitted pathway or joint Lasso.

### Pathway Selection

Figure 3 shows the estimated coefficients and p-values for two-stage Lasso-hierarchical Cox Model and two-stage hierarchical-hierarchical Cox Model (*s*_1_ = 1/*nλ*). Pathways with p-values less than 0.05 are labeled in Figure 3. We compared all significant pathways with core cancer pathways discussed in ENG et al. (2013) and listed those consistent core cancer pathways in Table 3.

**Figure 2.**
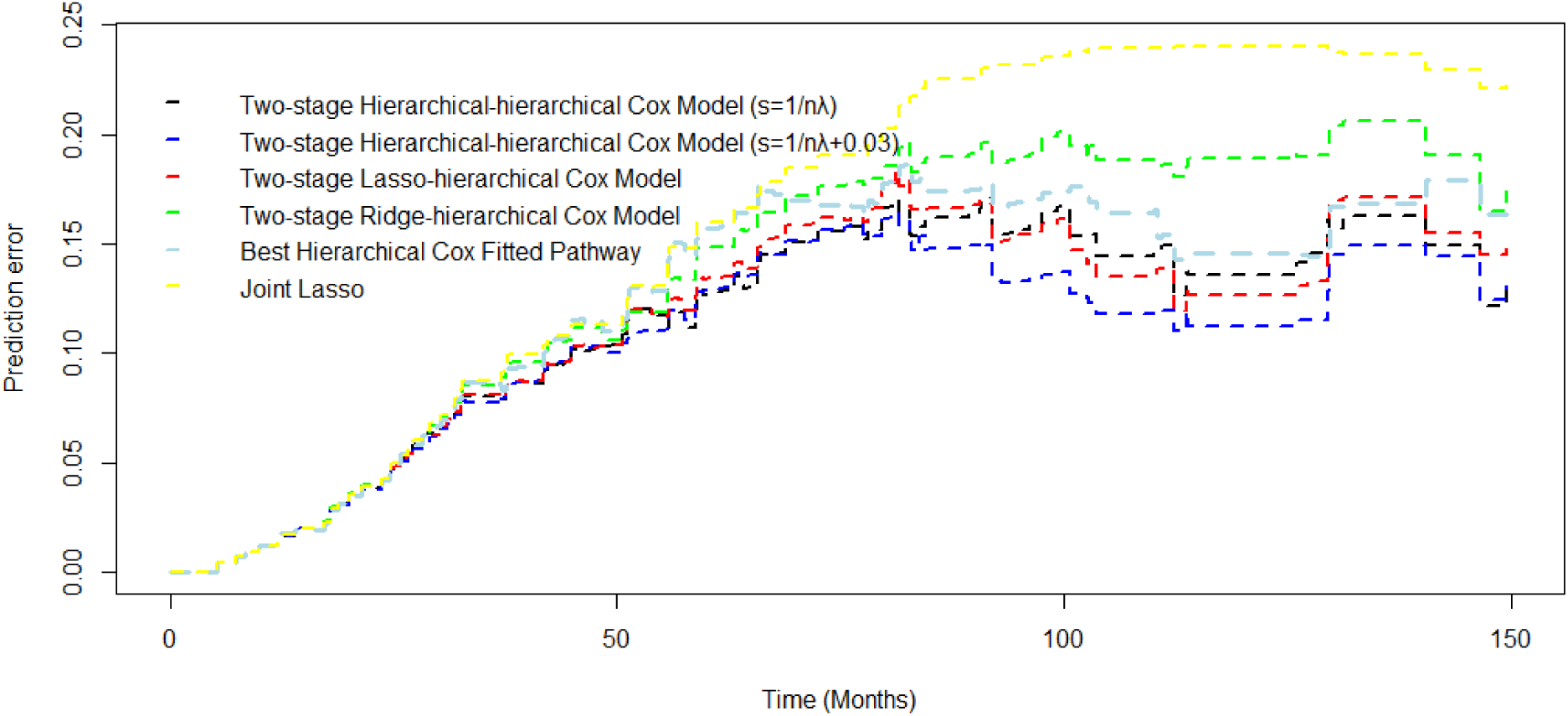
Brier prediction errors for Two-stage hierarchical-hierarchical Cox Model(*s*_1_ = 1/*nλ* and *s*_2_ = 1/*nλ* + 0.03), Two-stage Lasso-hierarchical Cox Model, Two-stage Ridge-hierarchical Cox Model, Best hierarchical Cox Fitted Pathway, and joint Lasso.

**Figure 3.**
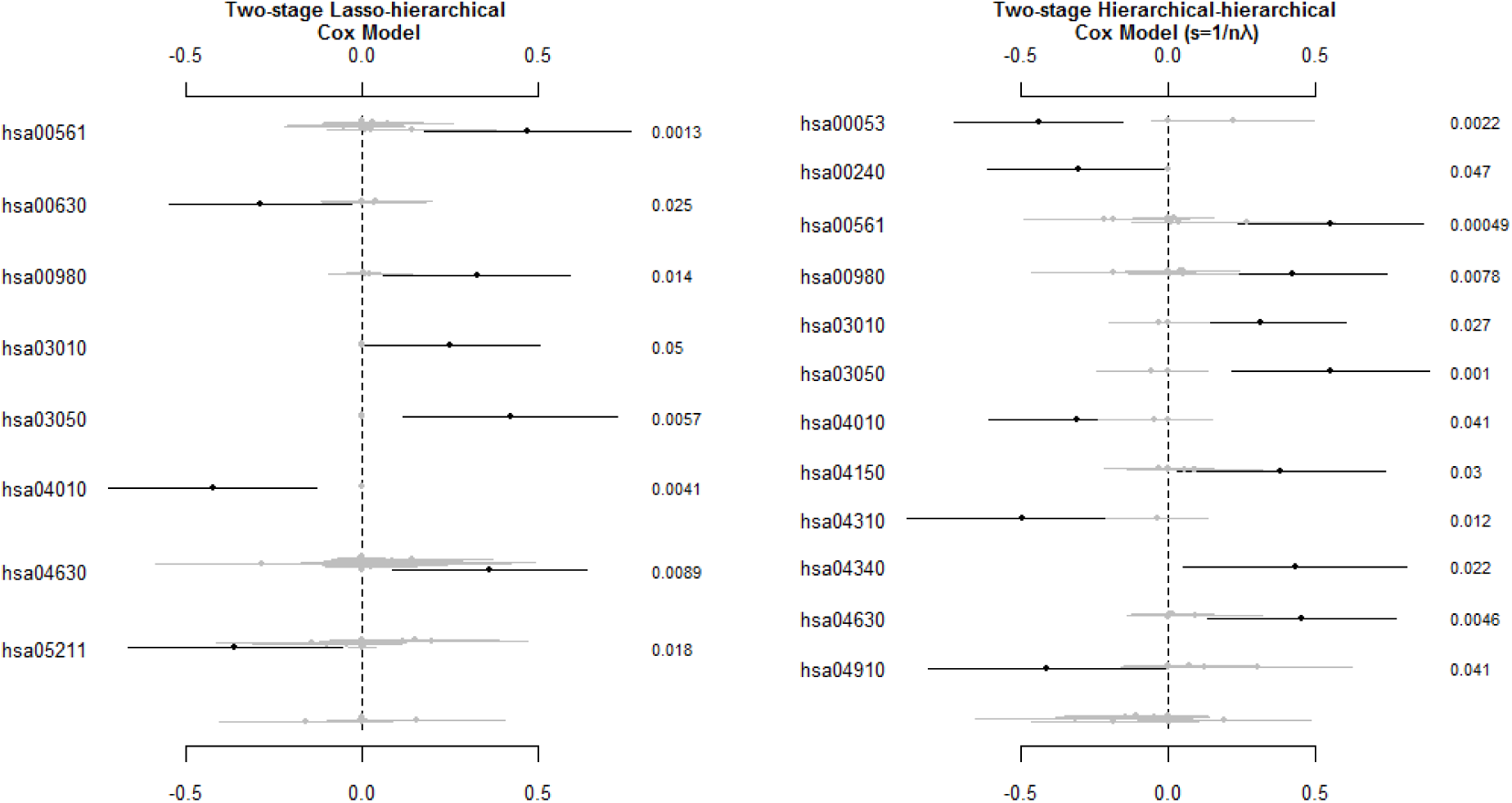
Estimated Coefficients and P-values of two-stage Lasso-hierarchical Cox model and two-stage hierarchical-hierarchical Cox model (*s*_1_ = 1/*nλ*), left column is the list of pathways; while the right column is the p-value for each pathway.

**Table 3.** Pathway selection comparison between two-stage Lasso-hierarchical Cox model and two-stage hierarchical-hierarchical Cox model (*s*_1_ = 1/*nλ*).

| Two-stage Lasso-hierarchical model | Two-stage Hierarchical-hierarchical model ($s_1 = 1/n\lambda$) |
| --- | --- |
| hsa04010:MAPK signaling pathway | hsa04010:MAPK signaling pathway |
| hsa04630:Jak-STAT signaling pathway | hsa04150:mTOR signaling pathway |
|  | hsa04340:Hedgehog signaling pathway |
|  | hsa04630:Jak-STAT signaling pathway |

### Risk Group Stratification

in order to demonstrate the potential for using two-stage approach to stratify patients into risk groups, we split the patients by the median of the cross-validated prognostic scores into two groups. Those patients with cross-validated prognostic score greater than the median were categorized as low-risk group; while patients with cross-validated prognostic score less than the median were categorized as high-risk group. The Kaplan-Meier curves for low-risk and high-risk groups from joint Lasso, joint hierarchical Cox model, two-stage Lasso-hierarchical (L-H) model and two-stage hierarchical-hierarchical (H-H) model (*s*_1_ = 1/*nλ*) are shown in Figure 4. Log-rank tests are carried out for each model. Both two-stage L-H and H-H models have significant differences of Kaplan Meier curves between low-risk group and high-risk group (p-value = 4.69e-11, p-value = 5.81e-11, respectively). Both joint Lasso and joint hierarchical Cox model have non-significant differences of Kaplan Meier curves between two groups (p-value = 0.242, p-value = 0.231, respectively).

**Figure 4.**
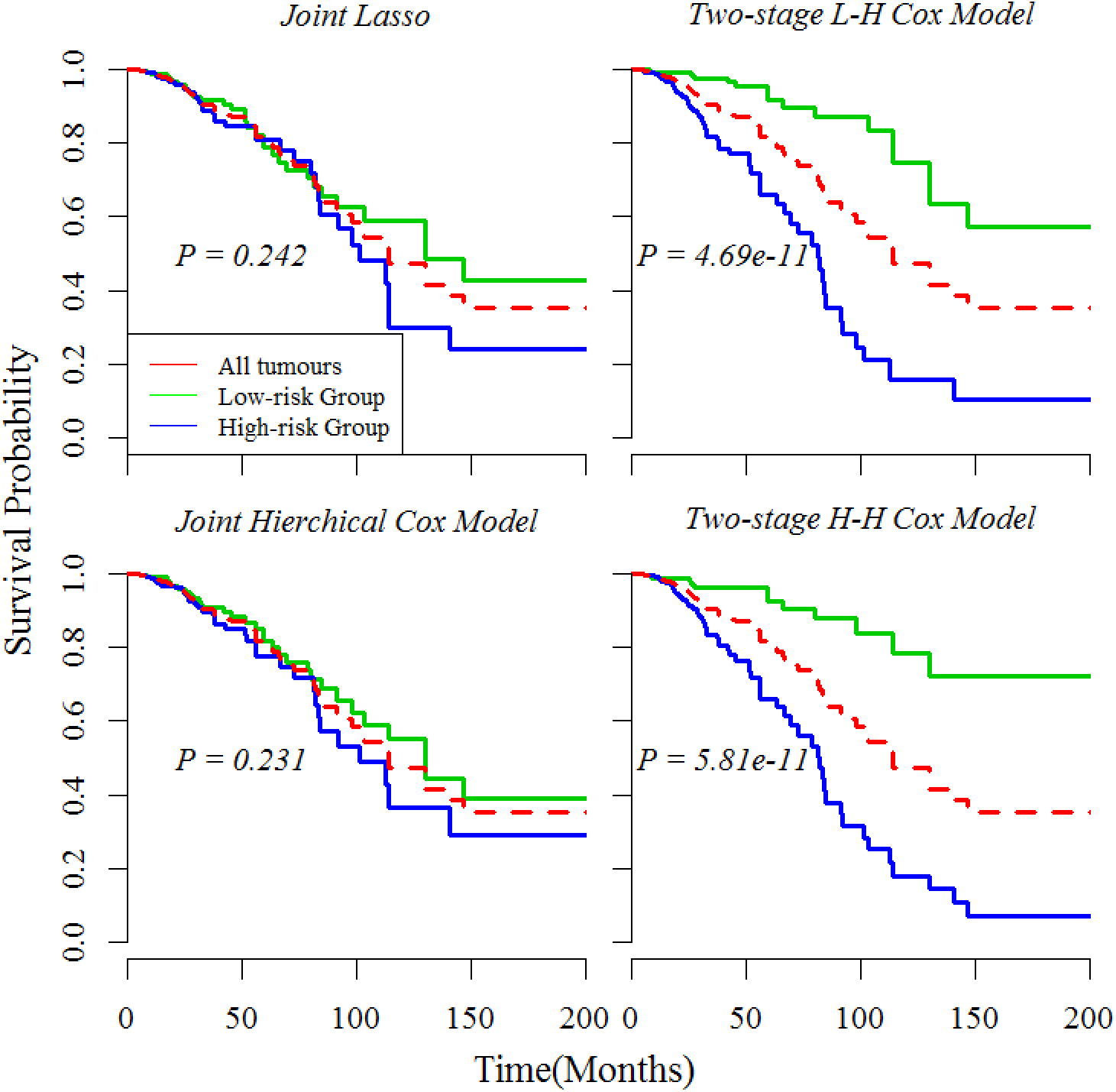
Kaplan-Meier Curves for low-risk and high-risk groups from joint Lasso, joint hierarchical Cox model, two-stage Lasso-hierarchical (L-H) model and two-stage hierarchical-hierarchical (H-H) model (*s*_1_ = 1/*nλ*), p-values are calculated using log-rank test.

## DISCUSSION

The heterogeneity of prognostic prediction in cancers has been a persisted problem for decades (BARILLOT 2013). It is now realized that cancer is a fundamentally disease of genome and can be understood by identifying the abnormal genes and proteins that are associated with the risk of developing cancer. Some statistical and machine learning methods have been used to analyze genomic data with gene-based approach to search for gene signature and to predict prognosis (RAPPAPORT 2007; BOVELSTAD *et al*. 2009; JACOB 2009; ZHANG *et al*. 2013; YUAN *et al*. 2014; ZHAO *et al*. 2014). Due to the complicated genetic nature of cancer and potentially underpowered statistical analysis of gene-based approach, it was suggested by Vogelstein that the complexity of cancer should be handled based on pathway-centric instead of gene-centric perspectives (JONES 2008). Oncogenes and tumor suppressor genes have been well studied and can be arranged into signaling pathways according to their biological functions such as cell survival, proliferation and metastatic dissemination. Other studies have investigated methods to analyze high-throughput cancer genomics data based on functional units, i.e. pathways (GOEMAN and BUHLMANN 2007; LEE *et al*. 2008; REYAL *et al*. 2008; ABRAHAM *et al*. 2010; TESCHENDORFF *et al*. 2010).

Our two-stage approach is developed to incorporate the functional structure of pathways to predict survival for cancer patients. Different from some previous methods that summarize pathway score using only gene information or positive/negative signs, our method incorporates the correlation with survival information in the calculating individual pathway information. Besides, we use cross-validated prognostic score as pathway score to be used in the second stage, which not only prevents overfitting and can be easily carried out, but also gives an unbiased view on the contribution of the different information from pathways to the prediction model. Our approach identified some important pathways, consistent with the core cancer pathways in ENG et al. (2013). Furthermore, all our two-stage approach pathway-based methods performed uniformly better than the gene-based joint models using Lasso or hierarchical cox model in terms of C-index, CVPL and reduced prediction error. Primarily, it can be seen that the highest C-index among the two-stage pathway-based model (0.760) is improved from the performance of both joint gene-based models (C-index = 0.507; 0.572 respectively) by around 0.2. Meanwhile, the Brier prediction errors of all two-stage models have been reduced from gene-based joint models greatly. The Kaplan-Meier analysis between low-risk and high-risk groups dichotomized by two-stage pathway-based models (p-value = 4.69e-11, p-value = 5.81e-11, respectively) have a remarkable performance in discriminating the prognostic effects between different patients compared with both joint gene-based models (p-value = 0.242, p-value = 0.231, respectively). Furthermore, among two-stage methods, the performance is dependent in the method selected to calculate the pathway score in the first stage. Lasso and hierarchical Cox models with the two estimated prior (*s*_1_ = 1/*nλ* and *s*_2_ = 1/*nλ* + 0.003) perform uniformly better than Ridge, Elastic Net and hierarchical Cox model with prior (*s*_3_ = 0.08).

Our approach is also capable of identifying core pathways in cancer. Mitogen-activated protein kinase (MAPK) pathways are important in controlling fundamental cellular processes, i.e. growth, proliferation, differentiation, migration and apoptosis (DHILLON *et al*. 2007). When abnormally activated, MAPK pathways can lead to the progression of cancer (USSAR and VOSS 2004; MCCUBREY *et al*. 2007). Another pathway, the mammalian target of rapamycin (mTOR), also plays an essential role in the regulation of cell proliferation, growth, differentiation, migration and survival. Similarly to MAPK pathways, the dysregulation of mTOR signaling happens in various human tumors, resulting in higher susceptibility to inhibitors of mTOR (HUANG and HOUGHTON 2003). The Hedgehog pathway regulates many fundamental processes including stem cell maintenance, cell differentiation, tissue polarity and cell proliferation. it has been demonstrated that inappropriate activation of Hedgehog pathway occurs in various cancers such as brain, gastrointestinal, lung, breast and prostate cancers (GUPTA *et al*. 2010). Furthermore, the JAK-STAT pathway is also identified in our approach. This pathway regulates in various cellular processes such as stem cell maintenance, apoptosis and the inflammatory response and was found frequently dysregulated in diverse types of cancer (THOMAS *et al*. 2015).

However, there are some potential limitations in this method. We implemented our approach only in microarray gene expression data from TCGA breast cancer project. There are many other platforms in gene expression data, such as RNA-Seq data. It may require additional constraints for the RNA-Seq data to be implemented into the model. Another potential limitation is that only 3181 mapped genes among the nearly 20000 expressed genes were fitted in the model, suggesting potentially loss of gene information.

Despite the above potential limitations, our two-stage pathway-based approach performs better in predicting overall survival of breast cancer and is able to identify important cancer pathways. In the future, we will apply our approach in other levels of genomic data, e.g. DNA methylation, miRNA and copy number alterations, for more than 30 types of cancer. We will also continue to develop more efficient ways to combine different levels of genomic data as well as clinical biomarkers into our proposed method to better predict cancer survival.

## Acknowledgments

This work was supported in part by the research grants: NIH 2 R01GM069430.

